# Positive selection in Europeans and East-Asians at the *ABCA12* gene

**DOI:** 10.1101/392811

**Authors:** Roberto Sirica, Marianna Buonaiuto, Valeria Petrella, Lucia Sticco, Donatella Tramontano, Dario Antonini, Caterina Missero, Ombretta Guardiola, Gennaro Andolfi, Heerman Kumar, Qasim Ayub, Yali Xue, Chris Tyler-Smith, Marco Salvemini, Giovanni D’Angelo, Vincenza Colonna

## Abstract

Natural selection acts on genetic variants by increasing the frequency of alleles responsible for a cellular function that is favorable in a certain environment. In a previous genome-wide scan for positive selection in contemporary humans, we identified a signal of positive selection in European and Asians at the genetic variant rs10180970. The variant is located in the second intron of the *ABCA12* gene, which is implicated in the lipid barrier formation and down-regulated by UVB radiation. We studied the signal of selection in the genomic region surrounding rs10180970 in a larger dataset that includes DNA sequences from ancient samples. We also investigated the functional consequences of gene expression of the alleles of rs10180970 and another genetic variant in its proximity in healthy volunteers exposed to similar UV radiation.

We confirmed the selection signal and refine its location that extends over 35 kb and includes the first intron, the first two exons and the transcription starting site of *ABCA12*. We found no obvious effect of rs10180970 alleles on *ABCA12* gene expression. We reconstructed the trajectory of the T allele over the last 80,000 years to discover that it was specific to *H. sapiens* and frequent among non-Africans already 45,000 years ago.

## Introduction

ATP binding cassette (ABC) transporters are trans-membrane ubiquitous proteins, that translocate natural substrates across plasma membranes. In humans, there are 49 genes coding for ABC transporters, arranged in eight subfamilies extensively studied because at least 11 of 49 genes are known to cause severe inherited diseases ^1^. The ATP-binding cassette, sub-family A member 12 (*ABCA12*) gene was discovered from cDNA of human placenta^2^. *ABCA12* is 207 kb long with fifty-three exons and two very long introns at its beginning (26.5 kb and 47.3 kb, respectively, Fig. 1A). *ABCA12* is a keratinocyte transmembrane transporter that binds and hydrolyzes ATP to transport lipids in the lamellar granules^3^. This activity is required to form the extracellular lipid barrier in the outermost layer of the skin, the *stratum corneum* of the epidermis^3^. The lipid barrier is composed of three lipid classes (cholesterol, free fatty acids, and ceramides) and acts as a primary barrier between the body and the environment to prevent excessive water loss and to avoid penetration of pathogens^4^. *ABCA12* has also a role in immunity: in macrophages it regulates the cellular cholesterol metabolism via an LXRb-dependent post-transcriptional mechanism^5^.

**Figure 1.**
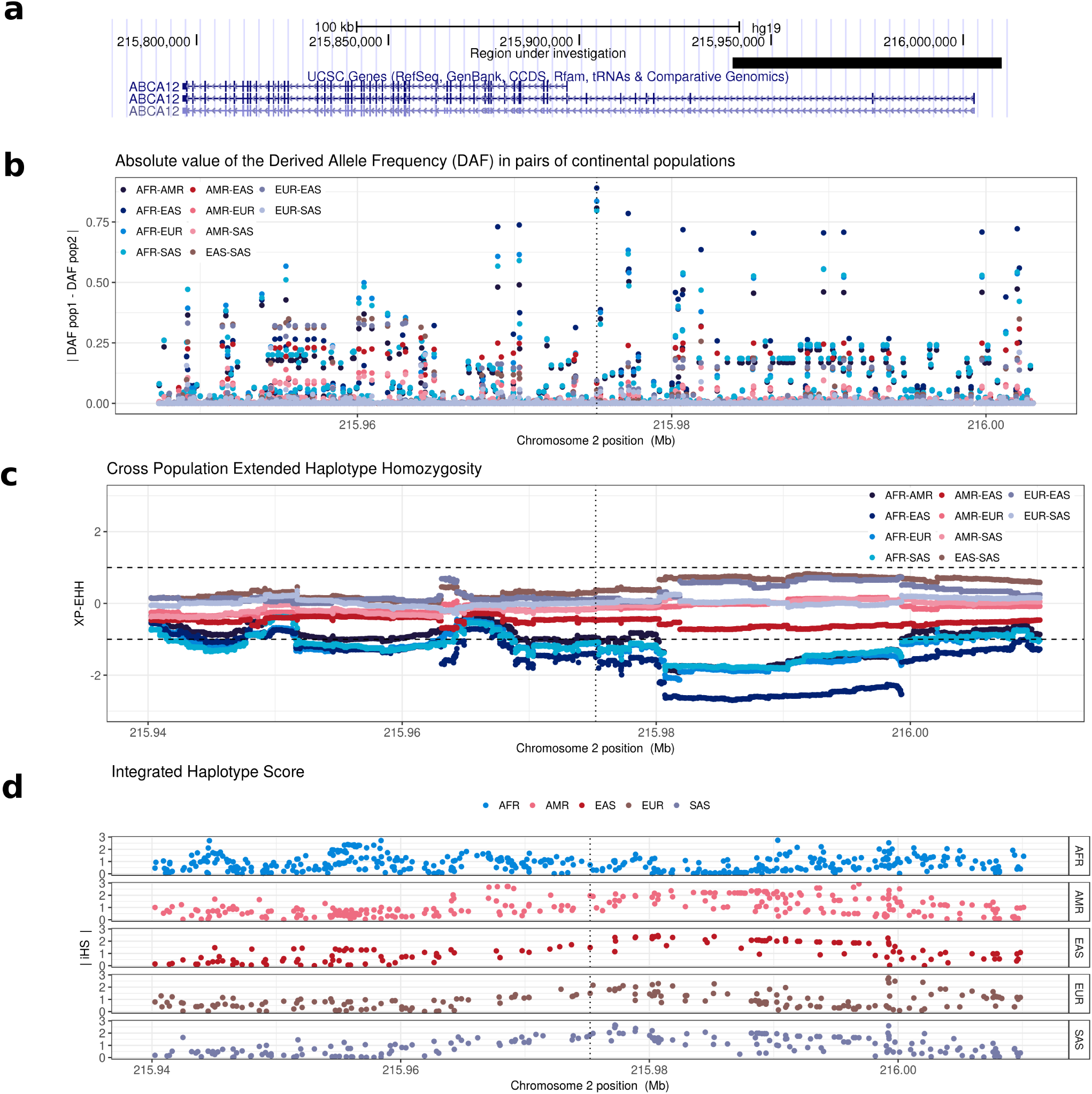
Positive selection at *ABCA12*. **a**) The *ABCA12* gene has is 55 exons and two long introns at its beginning. The black rectangle indicates the 70kb region surrounding rs10180970 considered in this project. (**b**) rs10180970 is the most differentiated variant between Africans and non-Africans as shown by the absolute difference of the derived allele frequency (∆DAF), however other variants seems to be also extremely differentiated. (**c**) The Cross Population Extended Haplotype Homozogysity statistic (XP-EHH), measured between pairs of continental populations, shows a signal of positive selection in non-Africans over 35kb downstream rs10180970, especially in East-Asians. (**d**) The signal is confirmed by the Integrated Haplotype Score (iHS) within continental populations. In panels **b**-**d** the dashed line indicates the genomic position of rs10180970

UV-radiation has a major effect on skin and keratinocytes and it is one of the most studied environmental stressors of the epidermal homeostasis. In keratinocytes, UV-radiation induces mutagenesis, apoptosis, proliferation, and metabolic changes. UV-radiation has a major role in determining skin pigmentation^6, 7^, but it also can have a detrimental effect on the lipid barrier reducing the intercellular lipid cohesion^8^. Earlier studies suggest that UVB radiation down-regulates epidermal *ABCA12* gene expression^9^ and in general UV light influences other ABC transporters activity in lymphocytes^10^.

The always growing availability of ancient DNA information allows to directly reconstruct the history of genetic mutations and understand if consequences of past adaptations are relevant for contemporary humans^11^. Past events of positive selection can be tested as detection of a major shift in allele frequencies^12^–^14^

In a genome-wide scan for positive selection in contemporary humans, we identified a signal of positive selection in Europeans and Asians at rs10180970 C/T, located in the second intron of *ABCA12*^15^. In the study, we demonstrated that the derived allele T is highly frequent in Asians and Europeans, compared to Africans, where the ancestral C variant prevails. This evidence, together with the identification of patterns typical of positive selection in the genomic region of rs10180970 let us speculate that ancestor of current Europeans and Asians should have had an advantage in carrying the T allele and that it would be interesting to determine the functional consequences and the trajectory of the T allele through time.

Here we present results from investigating the genomic region surrounding rs10180970. We confirm that this region underwent positive selection and investigate one possible selected phenotype. We have extended the set of individuals used to investigate genetic variation and natural selection at rs10180970, including also DNA sequences from ancient human samples. We confirm the signal of positive selection at rs10180970, investigate linkage with genetic variants in its proximity and the effect of the two alleles on *ABCA12* gene expression.

## Results

### Natural selection signal at the *ABCA12* gene extends 10kb downstream rs10180970

We identified rs10180970 as a possible candidate for positive selection in Europeans and Asians during a genome-wide scan for positive selection conducted on populations from Phase I 1000 Genomes Project^15^. In this study, we considered a 70kb region surrounding rs10180970 including the first two exons, the first and most of the second introns, and the transcription starting site of *ABCA12* (Fig. 1A). Compared to Phase I, the 1000 Genomes Phase III dataset^16^ includes 1,412 more samples belonging to 12 more populations (Supplementary Table S1) and in the region under study includes 1,612 variants for which the ancestral state is known and that we used to calculate statistics suggestive of population differentiation and positive selection.

Between pairs of continental populations, we evaluated the absolute difference of the derived allele frequency (∆DAF) and the Cross Population Extended Haplotype Homozygosity (XP-EHH^17^). ∆DAF approaches 1 when the derived allele prevails in one population and is almost absent in the other. According to ∆DAF, rs10180970 is still the most differentiated variant in comparisons of Africans and non-Africans in 70kb surrounding rs10180970 (Fig. 1 B). In Phase III the T/T genotype frequency in the combined sample of Europeans and Asians is significantly higher than Africans (Table 1, Fisher exact test p-value <2.2e-16). Similarly, in the Complete Genomics dataset (CG)^18^, the T/T genotype is significantly more frequent among Asians and Europeans compared to Africans. This is true both when considering samples partially overlapping with Phase I, and samples unique to CG (Table 1, Fisher exact test p-value <2.2e-16), suggesting that the observed pattern of higher derived allele frequencies in Europeans and Asians compared to Africans was not due to the specific set of sample used but is reproducible. XP-EHH detects recent positive selection highlighting selective sweeps in which the selected allele has approached or achieved fixation in one population but remains polymorphic in the overall population. In the 1000 Genomes Phase III dataset, we observe a stretch of negative XP-EHH outside the expected variance range (i.e., -1,1) in pairs of Africans with non-Africans. The signal is located approximately at 10kb downstream rs10180970, suggesting recent positive selection in non-Africans in this region (Fig. 1 C), and it is particularly strong in East-Asians. Within continental populations we calculated the Integrated Haplotype Score (iHS^19^). iHS detects the classic signal of strong directional selection through the identification of unusually long haplotypes with low diversity surrounding core SNPs. As for XP-EHH, also for iHS we observe peaks of unusually high iHS 10 kb downstream rs10180970 in non-African populations and especially in East-Asians (Fig. 1 D)

**Table 1.** Genotypes frequencies at rs10180970 in Phase III 1000 Genome and Complete Genomics datasets both full and non overlapping with 1000 Genomes Phase I (unique).

|  |  | T/T | T/C | C/C |
| --- | --- | --- | --- | --- |
| Complete Genomics<br>full dataset | Africans | 0 | 0.09 | 0.91 |
|  | Europeans & Asians | 0.89 | 0.10 | 0.01 |
| Complete Genomics<br>unique | Africans | 0 | 0.05 | 0.95 |
|  | Europeans & Asians | 0.88 | 0.11 | 0.01 |
| 1000 Genomes Phase III | Africans | 0.01 | 0.14 | 0.84 |
|  | Europeans & Asians | 0.85 | 0.14 | 0.01 |

Altogether these results show that while rs10180970 is the most differentiated variant in the *ABCA12* gene, haplotypes downstream rs10180970 differ between Africans and non-Africans. Haplotypes carrying the T allele of rs10180970 are very common in Europeans and Asians and tend to be longer and less diverse in these two populations compared to the others, suggesting that this region might have been affected by recent positive selection in these two populations.

Because linkage between variants in a long haplotype can be responsible for spurious signals, we computed linkage disequilibrium (LD) in continental populations between rs10180970 and 2,034 genetic variants in 70 kb surrounding rs10180970. For all populations but Africans, we observe LD within 10kb downstream rs10180970 (Figure 2A). In East-Asians and Europeans LD extends over 35kb downstream rs10180970, in a region inclusive of the first intron and the first exon. Within this block, LD is high in East-Asians (r^2^ around 0.75) and moderate in Europeans (r^2^ around 0.50). The genomic region in high LD contains functional features such as chromatin regulation (H3K4Me1 and H3K27Ac), enhancers and a promoter located in the proximity of the first intron and includes SNPs with ∆DAF>0.70 (Figure 2 B). Therefore we conclude that other SNPs are in high LD with rs10180970 in East-Asians and part of a long haplotype that is located in a genomic region with regulatory features.

**Figure 2.**
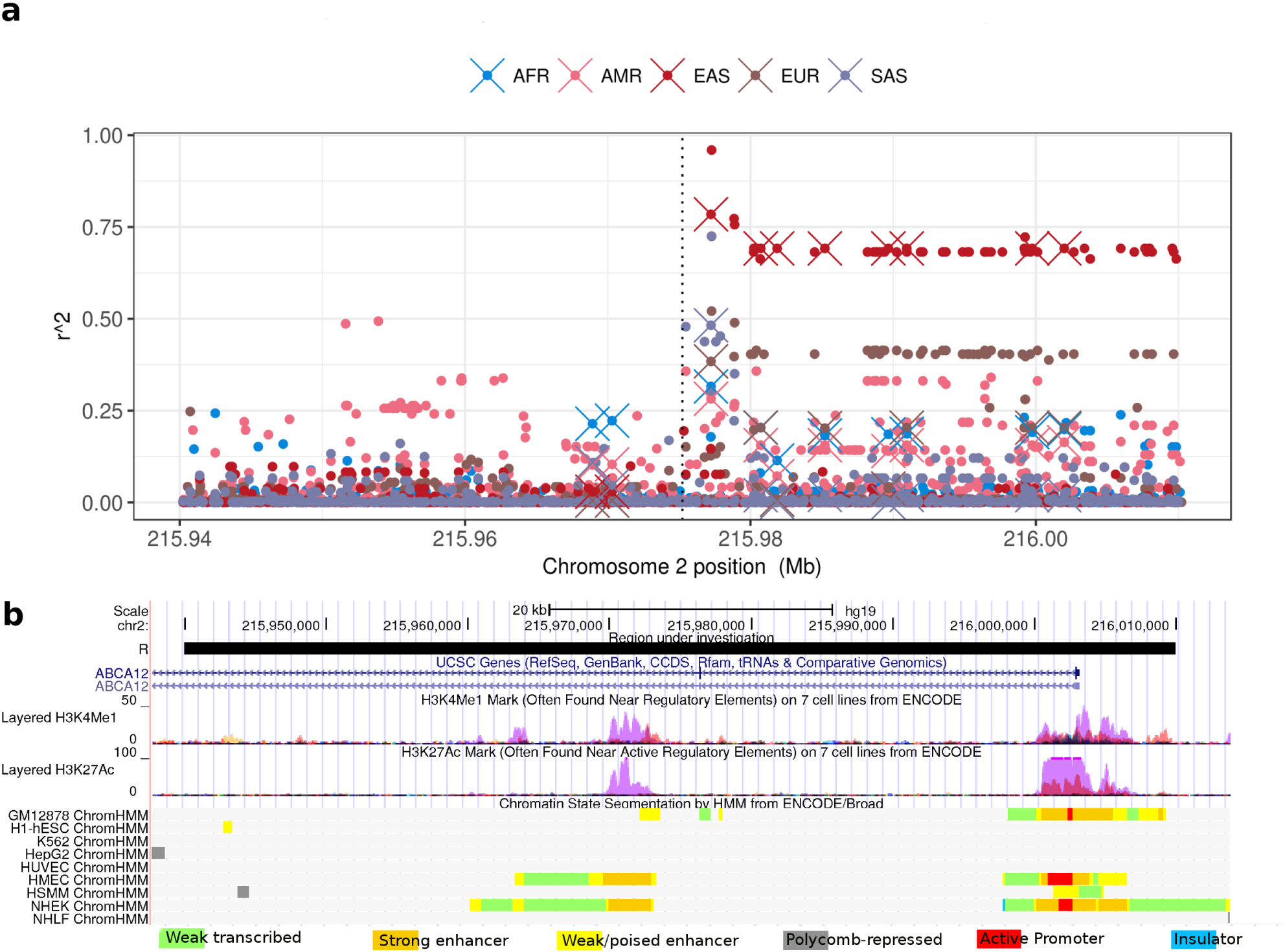
Linkage disequilibrium between rs10180970 and variants within 70 kb. (**a**) Vertical dotted line indicates the genomic position of rs10180970. Circles represent other variants at their genomic position on the x-axis and the r^2^ measures the non-random association of the alleles of each variant with rs10180970 on the y-axis. Crosses distinguish pairs in which the genetic variants have ∆DAF>0.70 as shown in Figure 1. (**b**) Screen-shot form Genome Browser showing chromatin modification and state segmentation for several cell types in the genomic region under investigation.

### No significant association between *ABCA12* expression and genotype at rs10180970

Having confirmed the signal of natural selection at rs10180970 we next asked if the two alleles have a different effect on gene expression. Furthermore, in the hypothesis that another variant in linkage with rs10180970 can be responsible for the functional changes, we decided to include also rs2970968 in the analysis. Besides being in linkage disequilibrium with rs10180970 in East-Asians and having high ∆DAF like other variants, rs2970968 has two further features that make it an interesting variant. First, we predicted the functional consequences of the variants with high ∆DAF and in linkage disequilibrium with rs10180970 in East-Asians using Funseq^20^ (Supplementary Table S2). We confirmed that rs2970968 (leftmost dark red cross in Figure 2A) is located in a region predicted to be an enhancer^21^ in lymphoblastoid, HeLa and HUVEC cell lines and that it is located in transcription factor binding peak of several transcription factors, although not in transcription factor motifs within peaks (Supplementary Figure S1). One of the transcription factors is implicated in mammalian skin tumors (ATF2^22, 23^) and three others are regulated by UVB radiation (STAT3, GATA3, and POU2F2)^24^–^26^. Secondly, rs2970968 is the closest SNP to a region that has shown to be essential for the functioning of the promoter of *ABCA12*^27^ (Supplementary Figure S1).

We investigated *ABCA12* expression in relation to the three genotypes of rs10180970 (C/C, C/T, T/T) and rs2970968 (A/A, A/G, G/G). Human skin would be the ideal tissue to test allele-specific gene expression, however sampling technique is invasive. It is instead possible to quantify *ABCA12* expression in total mRNA extracted from pulled hairs without the need for invasive skin biopsies^28^. Because *ABCA12* expression is influenced by sun exposure^29^, to discriminate between the genetic and the environmental effects, we performed the experiment in a group of individuals of different geographical origin but all resident in the same city. We determined the genotypes at rs10180970 and rs2970968 through DNA sequencing and measured *ABCA12* expression through quantitative PCR of reverse transcribed total mRNA extracted from pulled hair roots. We considered as reference the average *ABCA12* gene expression of genotypes of the ancestral alleles (C/C for rs10180970, and AA for rs2970968), which are also the allele prevalent in Africans, and calculated the fold change of the other observed genotypes. After quality controls, we had data on both genotype and *ABCA12* expression for 36 and 39 samples at rs10180970 and rs2970968, respectively. As shown in Figure 3, there is no significant correlation between *ABCA12* gene expression and genotypes at both loci (One-way non-parametric ANOVA p-values>0.9 for both sites), suggesting no effect of rs10180970 or rs2970968 genotypes on *ABCA12* expression in hair roots. Therefore, we conclude that the effect of the selection at these two loci on gene expression is not obvious when individuals are exposed to similar UV radiation.

**Figure 3.**
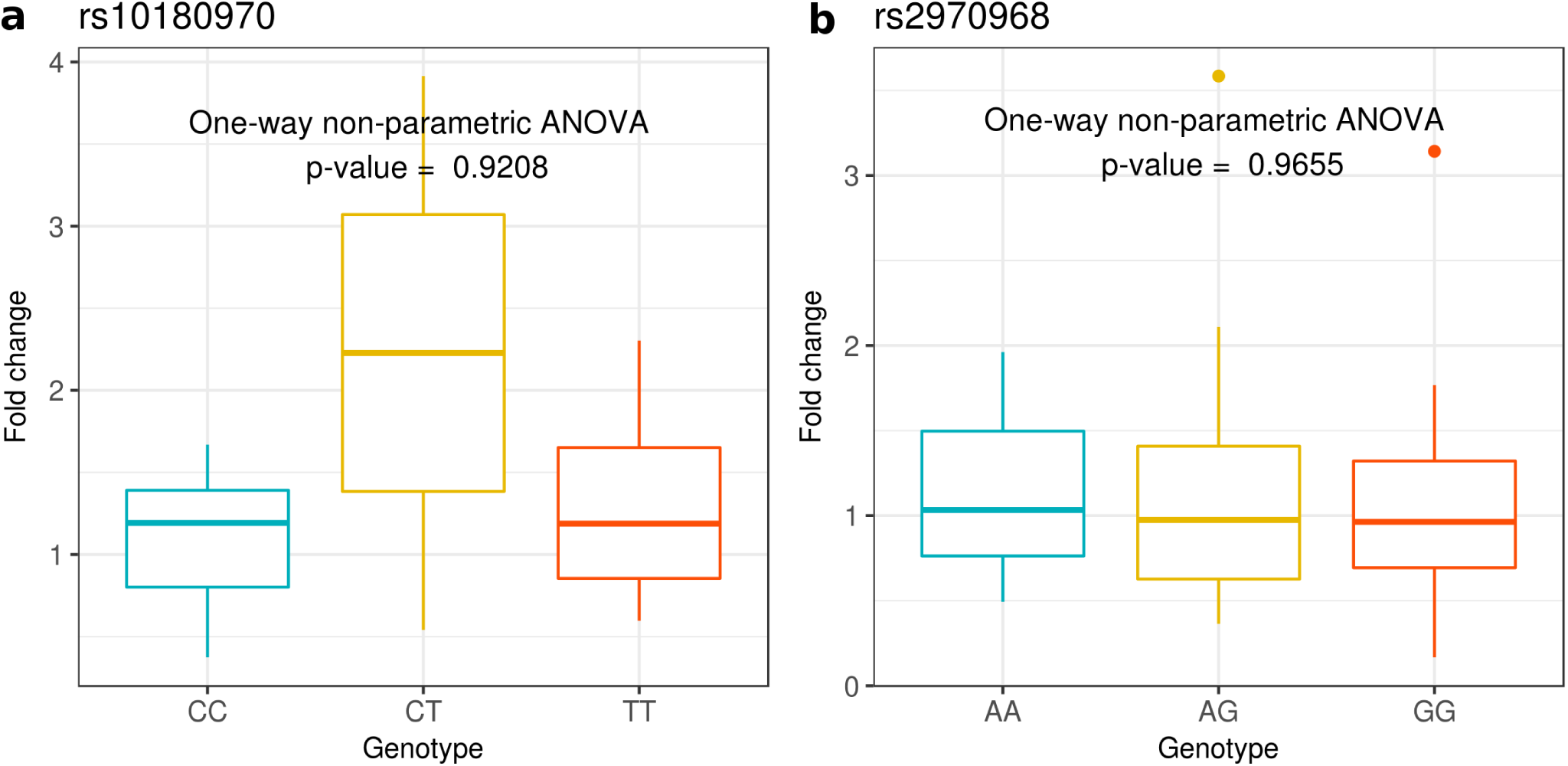
*ABCA12* expression in relation to the three genotypes of (a) rs10180970 and (b) rs2970968. Fold change refers to average *ABCA12* gene expression of ancestral genotypes, which are also the genotypes prevalent in Africans (C/C for rs10180970, and AA for rs2970968).

### The T variant of rs10180970 is specific to *H. sapiens* and rapidly increased in frequency in non-Africans

We determined the pattern of genetic variation at rs10180970 in time and space in 88 ancient and 2,764 contemporary samples ranging the last 80,000 years with the aim to understand when the T allele first appeared and how its frequency evolved through time. Among contemporary samples, 41 were collected for this study and are from 9 geographically diverse populations (Supplementary Table S1). Their genotype at rs10180970 was determined by Sanger sequencing a 600 bp region containing the variant. For the rest of the contemporary samples genotypes at rs10180970 were publicly available from the 1000 Genomes Phase III^16^ and a study on West Africans^30^. For ancient samples, we used publicly available raw sequencing data (Supplementary Table S3) to perform variant calling. Beside *H. sapiens*, ancient samples include also one *Denisova* sample and two *Neanderthals*.

The allele frequency of the derived allele T (*f* _T_) through time was determined grouping individuals in bins of 2000 years (Fig.4). Some pools are made of very few individuals (n=1 in extreme cases). Individuals were subdivided according to species and within *H. sapiens* in Africans and non-Africans. In our data, the T allele appeared for the first time around 45 kya in a *H. sapiens* sample from Russia^31^ and rapidly *f* _T_ reached 0.9 in non-Africans remaining stable until our days.

Of the 16 African ancient samples available to date, only three have reliable data in the rs10180970 region (Supplementary Table S3). The most ancient African data is from Ethiopia 4.5kya^32^, while the other two are from 1.3-2kya^33^. All ancient Africans have genotype C/C. In contemporary Africans *f* _T_ is very low but not completely absent(0.09), while in North-Africans *f* _T_ is on average 0.4 and T is the major allele in Egyptians, within North-Africans (Fig.4 B). Admixture analysis shows that the Africans and North-Africans with higher *f* _T_, in fact, have quite high European and South Asian components probably reflecting recent admixture (Fig.4 C.

Both Neanderthals and the one Denisova sample available are homozygous for the ancestral C allele, suggesting that the T allele might have been specific to *H. sapiens*, as far as we know about the other two species. In support of this hypothesis, two recent researches find that there is a very little probability of admixture in the region surrounding rs10180970 between *H. sapiens* and Neanderthals^34^ (Supplementary Figure S2) and *H. sapiens* and Denisova^35^.

We conclude that the T variant of rs10180970 is specific to *H. sapiens* and rapidly increased in frequency in non-Africans, while the low prevalence of T in contemporary Africans might be due to recent admixture with Europeans and South-Asians.

## Discussion

Humans are exposed to environmental agents through the naked skin, and in fact, strong natural selection has been detected in a number of skin-related phenotypic traits such as hair and eye pigmentation^6, 36, 37^, and hair thickness^38, 39^. The lipid barrier is the most external layer of the skin and it acts as an interface with the environment, providing protection against UV, dehydration, and pathogens. The formation of lipid barrier is a complex process and mutations in genes involved in this process result in skin diseases. Nevertheless, to date, to our knowledge, no studies investigated natural selection of genes that contribute to build the lipid barrier such as *ABCA12*.

The genetic variant rs10180970 located on chromosome 2 in the *ABCA12* gene, is predicted to be under positive selection in Europeans and Asians, as we discovered in a genome-wide scan based on population differentiation^15^. Here we investigated the genomic region containing rs10180970 to ask whether the variant is truly under positive selection and investigated the functional consequences of the selection event.

We confirmed the signal of selection in Europeans and Asians and discovered high linkage disequilibrium in a 35kb region downstream rs10180970 (Fig. 2). In the 35kb region the signals of positive selection are as strong or stronger than those initially found at rs10180970 (Fig. 1) and include the promoter of the gene (Fig. 2). By using a larger dataset of DNA sequences and a more fine population subdivision compared to the one used to discover the signal, we observe that the signal is stronger in East-Asians compared to South-Asians, possibly because South-Asians exposure to UV radiation is more similar to that of Africans. Despite confirming the positive selection signal we could not observe obvious allele-specific effect of rs10180970 on *ABCA12* gene expression (Fig. 3) in a sample of individuals exposed to the same level of UV radiation. Finally, we traced the history through time of the allele under selection using publicly available ancient human sequences and established that the mutation is specific to *H. sapiens* (Fig. 4, Supplementary Figure S2). While our findings strongly corroborate the signal of selection they also suggest that a different functional approach should be used to decipher the functional consequences of the event of selection in terms of gene expression.

**Figure 4.**
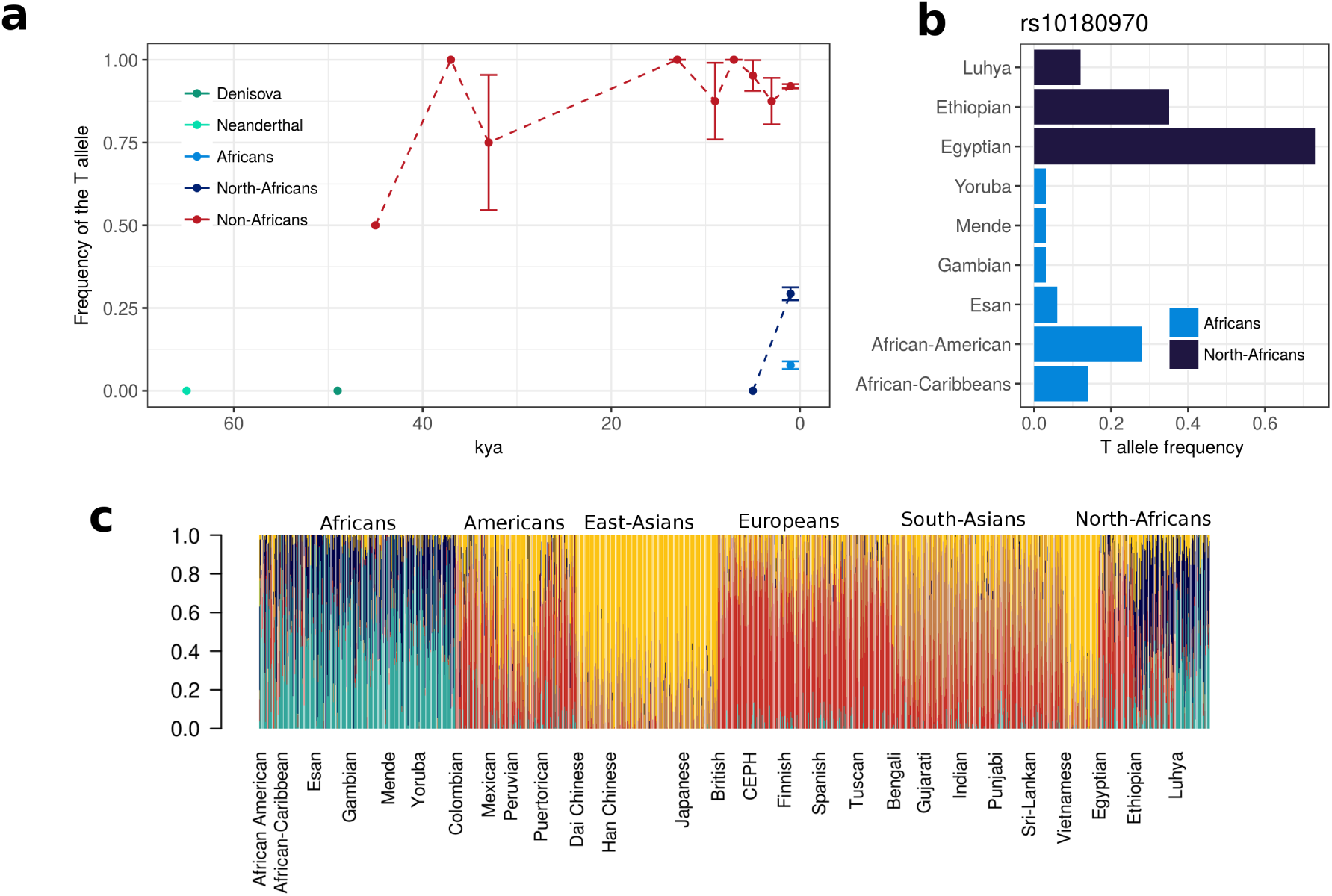
Prevalence of the T allele of rs10180970 in time and space. (**a**) T allele trajectory in the last 80,000 years in 88 ancient and 2,764 contemporary samples. The T allele is observed for the first time 45,000 years ago in non-Africans and rapidly rose in frequency. (**b**) T allele frequency is very low among African populations with the exception of North-Africans (especially Egyptians and Ethiopians) and African American. (c) T allele was probably recently introgressed in African American, as admixture analysis show a shared genetic component of African American with Europeans and East-Asians in the region under investigation. Vertical bars represent single individuals and colors reflect membership to admixture clusters in the most likely hypothesis of five clusters.

We suggest that further work is required to fully decipher the effect of selection on *ABCA12*. While it is not possible to model the gene effect in mouse because there is no homologous region in mouse, we found that *ABCA12* is well expressed in HaCaT cells, a spontaneously transformed aneuploid immortal keratinocyte cell line from adult human skin^40^. As a preliminary work, we silenced *ABCA12* in HaCatT cells using small interfering RNAs and confirmed that *ABCA12* expression is reduced after UVB radiation (20 mJ and 30 mJ), with downregulation remaining sustained up to 18 hours after treatment (Supplementary Figure S3). We also determined by Sanger sequencing that HaCaT cells are heterozygous at rs10180970 and therefore can be used for allele-specific expression assessment at this locus. On the other hand, because of the possibility that other sites in linkage with rs100180970 are involved, it would be probably worth to consider sequence information at other sites as well. More in general, overall indication for future research implicate the study of the non-coding regions of *ABCA12*, like sequence/haplotype-specific enhancer assay, even if the dimensions of the gene can be a limitation.

There are two main findings from our work. First, we reconstructed the demographic history of the T allele using the currently available information to discover that it is specific to *H. sapiens* and it was apparently very frequent in humans that migrated out of Africa prior to 50kya. The second main finding is that while rs10180970 might act as a sentinel for the signal of selection, the signal extends over 35 kb and includes the first intron, the first two exons and the transcription starting site of *ABCA12*. Linkage disequilibrium in this region is high in East-Asians and Europeans, there are several regions predicted to be enhancers and two regions predicted to be sites of chromatin modifications (methylation and acetylation), thus possibly influencing gene expression.

Our study endorses and refines previous studies. Similar to *ABCA12*, other members of the ABC family have also been identified in previous scans for positive selection^41^. The genomic region containing *ABCA12* was identified as one of the most differentiated between continental populations^42^, and in fact *ABCA12* hosts one of the top ten SNPs in the genome, useful to discriminate continental populations in clustering analyses^43^. *ABCA12* was identified as one of the top candidate genes in a study investigating local adaptation in Asians^44^ and stands out in a selection scan performed with LD-based statistics on HapMap Phase II data^45^. However, none of these studies provides an indication of the exact genomic location of the signal within the gene. In this study, we corroborate the signal of selection at rs10180970 that was localized in our previous study^15^, and provide initial evidence that will help understand why the T allele of rs10180970 was favored in the ancestors of current Asians, in particular, East-Asians, and Europeans. Our work also shows that with the exception of few cases (e.g.^46, 47^) it is surprisingly difficult to identify the selected allele and selected phenotype, even with abundant sequence data and functional tests.

## Methods

### Samples and genotyping

Thirty-Nine samples were collected from healthy fully informed, consenting, adult volunteers from 9 countries (Supplementary Table S1). Saliva samples were collected using the Oragene DNA (OG-500) kit. Genomic DNA was phenol–chloroform extracted from saliva^48^. The study was approved by the ethics committee of the University of Naples Federico II (protocol number 106/15) and conducted in accordance with the relevant guidelines and regulations. Informed consent was obtained from all participants.

Genotypes at rs10180970 and rs2970968 were obtained by Sanger-sequencing a 325bp and 400bp regions, respectively, of the *ABCA12* gene where the variants are located. The fragments were amplified from genomic DNA by PCR in a final volume of 50µl using oligos designed with Primer3^49, 50^ and the following conditions: 1 cycle 95°C for 5’; 35 cycles at 95°C for 1’, 62°C for 1’, 72°C for 45′; 1 cycle at 72°C for 10’. PCR products were purified using PCR purification kit from QIAGEN.

All other contemporary samples in the study are part of public collections and genotype data at rs10180970 was available in form of vcf files ^16, 18, 30^. For ancient samples we screened publicly available sequence reads data of 400 samples spanning 80,000-200 years before present^31, 51–64^, however filtering for coverage >2 in the region of interest resulted in 88 samples listed in Supplementary Table S3. Variant calling in ancient samples was performed with samtools/bcftools^65^.

#### Derived allele frequencies, linkage disequilibrium, admixture, introgression, natural selection analyses

Derived allele frequencies were computed using vcftools^66^ using the information on derived allele from the 1000 Genomes Project^16^. Linkage disequilibrium (LD) between rs10180970 and any other SNP in ±2Mb was computed with PLINK1.9^67^. Clustering analysis in a region of 2Mb surrounding rs10180970 including 55,683 Single Nucleotide Polymorphisms (SNPs) was done with ADMIXTURE^68^. ADMIXTURE was run under the hypotheses of 2 to 10 clusters with cross-validation computation.

Data for checking if the region of rs10180970 is among the ones putatively introgressed form Neanderthal were downloaded from http://genetics.med.harvard.edu/reichlab/Reich_Lab/Datasets_-_Neandertal_Introgression.html.

Integrated haplotype score (iHS) and Cross-Population Extended Haplotype Homozygosity (XP-EHH) were calculated with Hapbin^69^. Prameters for iHS calculation were –minmaf 0.01, –cutoff 0.01. iHS and XP-EHH log-ratio were normalized to have zero mean and unit variance using the standard normalization implemented in Hapbin that groups variants in bins of 2% frequency.

#### RNA extraction and processing from hair follicles

Hair follicles (5 to 7 bulbs) were grasped as near to the scalp as possible without damaging hair roots. Total RNA was extracted from bulbs on the day of collection, or following storage at 4°C for 1–5 days in RNALater and placed into a 1,5 ml microcentrifuge tube containing 1 ml of Trizol Reagent (Fisher Molecular Biology). RNA extraction was then conducted according to manufacturer’s instructions. 500 ng RNA were reverse-transcribed using SuperScript^®^ VILO™ MasterMix (Invitrogen). qPCR was performed on technical triplicates per each sample using Light Cycler 480 ^®^ II (Roche), on 2 µl of previously diluted cDNA (1:5) template using LightCycler ^®^ 480 SYBR Green I Master Mix (Roche), according to the manufacturer instructions. The thermal profile consisted of 1 cycle at 95°C for 10’; 40 cycles at 95°C for 10′, 60°C for 10′, 72°C for 10′. Fold changes were calculated relative to Hypoxanthine Phospho Ribosyl Transferase (*HPRT1*) expression as in^70^

#### HaCat cells culturing, genotyping, silencing and UV radiation

HaCat cells were cultured in 75 cm^2^ tissue culture flasks in Dulbecco’s Modified Eagle’s Medium, containing 10% fetal bovine serum and 1% penicillin and streptomycin. Cells were maintained in a humidified atmosphere (5% CO2) at 37°C. Genomic DNA was extracted from HaCat cells^71^ and genotype at rs10180970 was determined by Sanger sequencing.

For silencing experiments, HaCaT cells were transfected with a *ABCA12*-targeting siRna (Sigma Aldrich). Transfection was performed with Oligofectamine following the manufacturer’s protocol. Cells were collected 72 hours after transfection. Silencing efficiency was evaluated by Real Time Quantitative PCR (RT-qPCR) following total RNA extraction using RNeasy Mini Kit-QIAGEN and 1µg and reverse-transcription using SuperScript^®^ VILO™ MasterMix (Invitrogen).

For UVB radiation HaCaT cells were plated in 60mm plates and grown to 70-90% confluence. Subsequently, cells were irradiated with 20 and 30 mJ/cm^2^ using three lamps (Philips Ultraviolet 8 TL 20W/01 RS lamps; Philips, Eindhoven, Netherlands) generating UVB light in the range of 290–320 nm with an emission peak at 312 nm. The intensity of UVB irradiation was measured using a phototherapy radiometer (International Light, Newburyport, MA) of UVB. After irradiation cells were incubated for 0,6 and 18 hours and *ABCA12* expression was assessed by Real Time PCR (RT-PCR). Fold changes were calculated relative to Ribosomal Protein Large P0 (*RPLP0*) expression as in^70^.

## Data Availability

The datasets generated during and analysed during the current study are available from the corresponding author on reasonable request.

## Acknowledgements

We would like to thank all the anonymous volunteers that participated in the project. We also thank Cristina D’Aniello for helping with DNA extraction protocol and Erik Garrison for assisting sampling. A special thanks to Lucia De Martino for helping to submit the project to the ethical committee. C.TS., Q.A, and Y.X. were supported by the Wellcome Trust grant 098051.

## Author contributions

V.C., C.M., Q.A., C.T.S., M.S., G.D.A. conceived the experiment(s). V.C., R.S., M.B., V.P., L.S., D.T., D.A., O.G., G.A. conducted the experiments. V.C., R.S., M.S., G.D.A., analysed the results. V.C., R.S., M.B., Y.X., Q.A., C.T.S., M.S., G.D.A. worked at the interpretation of the results. V.C., M.S., G.D.A. wrote the manuscript. All authors reviewed the manuscript.

## Supplementary Information

**Table S1.**
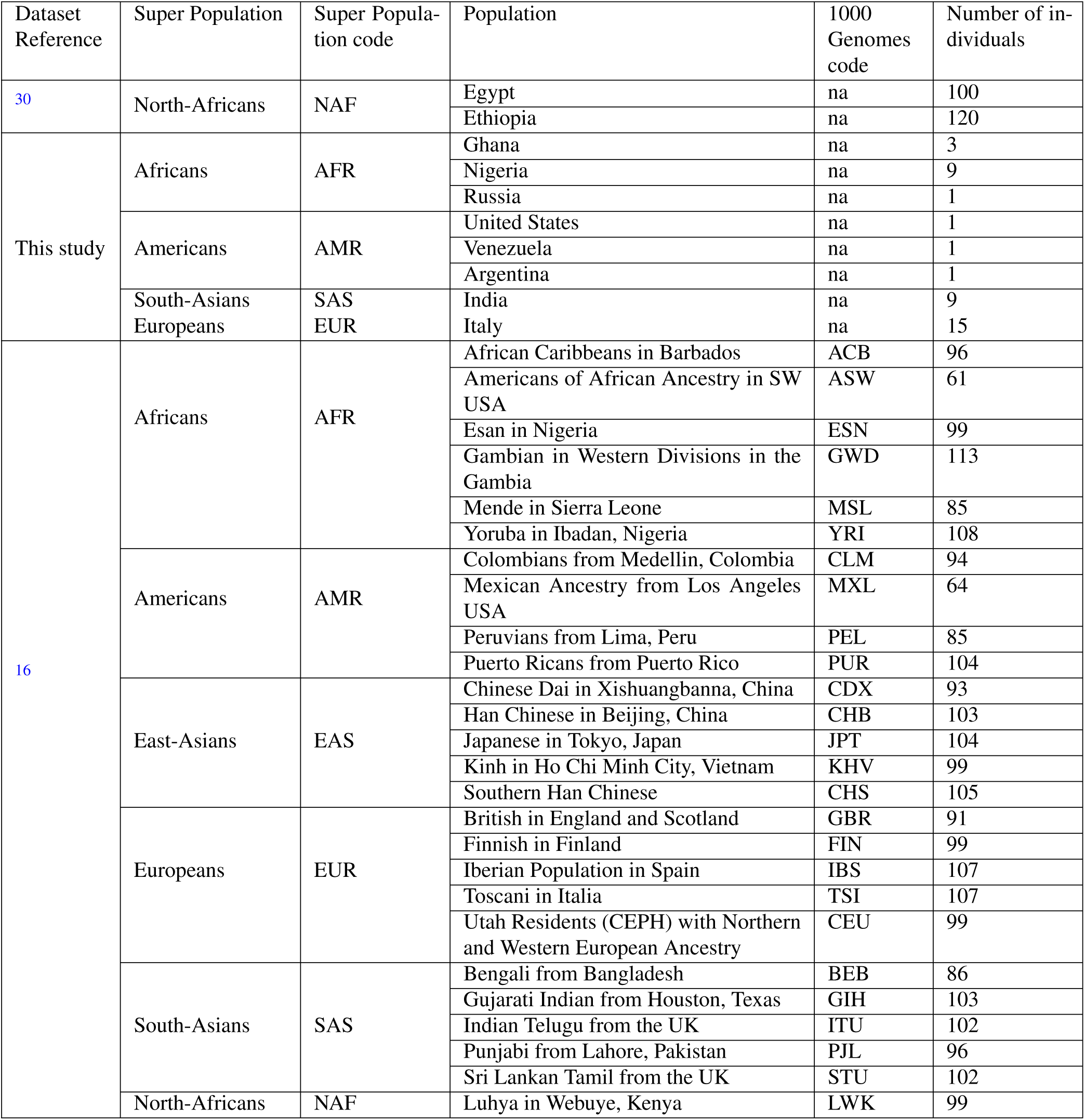
Geographical origin of the samples in this study.

**Table S2.** Results of the functional analysis using Funseq^20^, of the variants with high ΔDAF and in linkage disequilibrium with rs10180970 in East-Asians.

| Variant | Coding | ENCODE annotation | Motif breaking | Sensitive | Ultra sensitive | Non-coding score |
| --- | --- | --- | --- | --- | --- | --- |
| hline<br>rs2948974 | No | . | . | . | . | 0 |
| rs2948975 | No | TFP(STAT3\|chr2:215970293-215970991) | . | . | . | 1 |
| rs10180970 | No | . | . | . | . | 0 |
| rs60395874 | No | . | . | . | . | 0 |
| rs10206315 | No | . | . | . | . | 0 |
| rs35127007 | No | . | . | . | . | 0 |
| rs34010652 | No | . | . | . | . | 0 |
| rs10165506 | No | . | . | . | . | 0 |
| rs10182390 | No | . | . | . | . | 0 |
| rs2970966 | No | . | . | . | . | 0 |
| rs2970968 | No | DHS (MCV-27 \| chr2:216001900-216002050), Enhancer (chmm/segway \| chr2:216001000-216003891), TFP (EBF1 \| chr2:216001332-216002896), TFP (EP300 \| chr2:216001847-216003037), TFP (POU2F2 \| chr2:216001396-216002207), TFP (STAT3 \| chr2:216001267-216002838), TFP (STAT3 \| chr2:216001305-216002967), TFP (STAT3 \| chr2:216001343-216002951), TFP (STAT3 \| chr2:216001959-216002946 ) | . | . | . | 1 |

**Table S3.** Summary of the data used to call variants from ancient samples ordered from the most ancient sample.

| Reference | n samples | Species | Age (kya) |
| --- | --- | --- | --- |
| Prufer_2014<sup>51</sup> | 2 | Neanderthal | 65-80 |
| Meyer_2012<sup>52</sup> | 1 | Denisova | 50 |
| Gunther_2015<sup>53</sup> | 1 | *H. sapiens* | 48 |
| Fu_2015<sup>31</sup> | 1 | *H. sapiens* | 45 |
| Seguin_Orlando_2014<sup>54</sup> | 1 | *H. sapiens* | 37 |
| Raghavan_2014<sup>55</sup> | 3 | *H. sapiens* | 24 |
| Jones_2015<sup>56</sup> | 2 | *H. sapiens* | 13-5.8 |
| Rasmussen_2014<sup>57</sup> | 1 | *H. sapiens* | 10.7 |
| Lazaridis_2014<sup>58</sup> | 2 | *H. sapiens* | 7.5-5.8 |
| Haak_2015<sup>59</sup> | 31 | *H. sapiens* | 6.3-1 |
| Skoglund_2014<sup>60</sup> | 3 | *H. sapiens* | 4.8 |
| Skoglund_2017<sup>33</sup> | 3 | *H. sapiens* | 3-2.5 |
| Sikora_2017<sup>72</sup> | 5 | *H. sapiens* | 34 |
| Allentoft_2015<sup>61</sup> | 26 | *H. sapiens* | 4.2-1.1 |
| Rasmussen_2010<sup>62</sup> | 1 | *H. sapiens* | 4 |
| Malaspinas_2014<sup>63</sup> | 2 | *H. sapiens* | 0.3 |
| Raghavan_2015<sup>64</sup> | 4 | *H. sapiens* | 0.2 |

**Figure S1.**
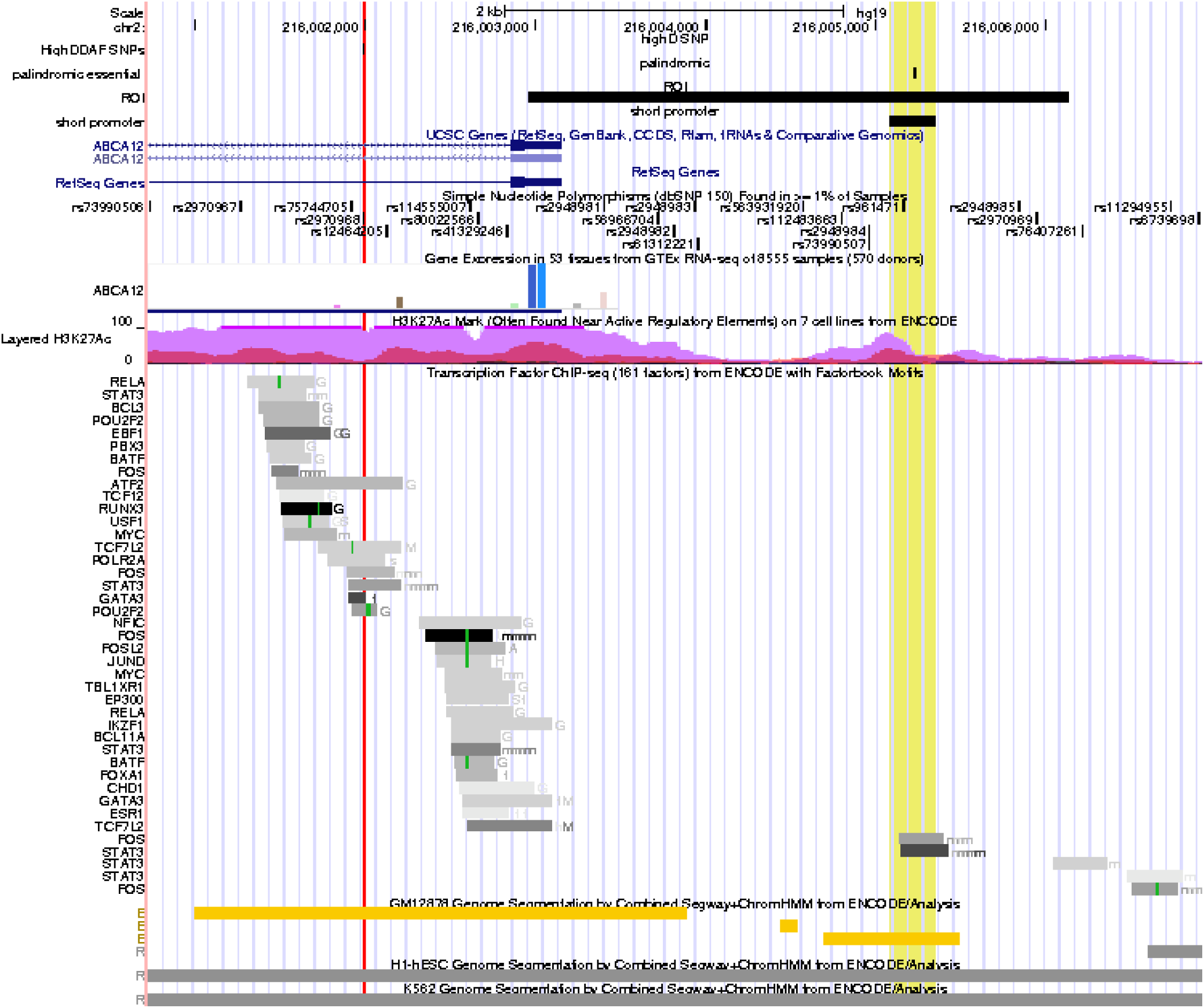
Genomic region surrounding rs2970968. The red vertical line indicates the genomic location of rs2970968 in a region predicted to be an enhancer in lymphoblastoid, HeLa and HUVEC cell line. rs2970968 is also located in transcription factor binding peak of several transcription factors of which one is implicated in mammalian skin tumors (ATF2) and four are regulated by UVB radiation (STAT3, GATA3, and 2/19POU2F2). It is also the closest SNP to a region that has shown to be essential for the functioning of the promoter of *ABCA12*, highlighted in yellow in the figure.

**Figure S2.**
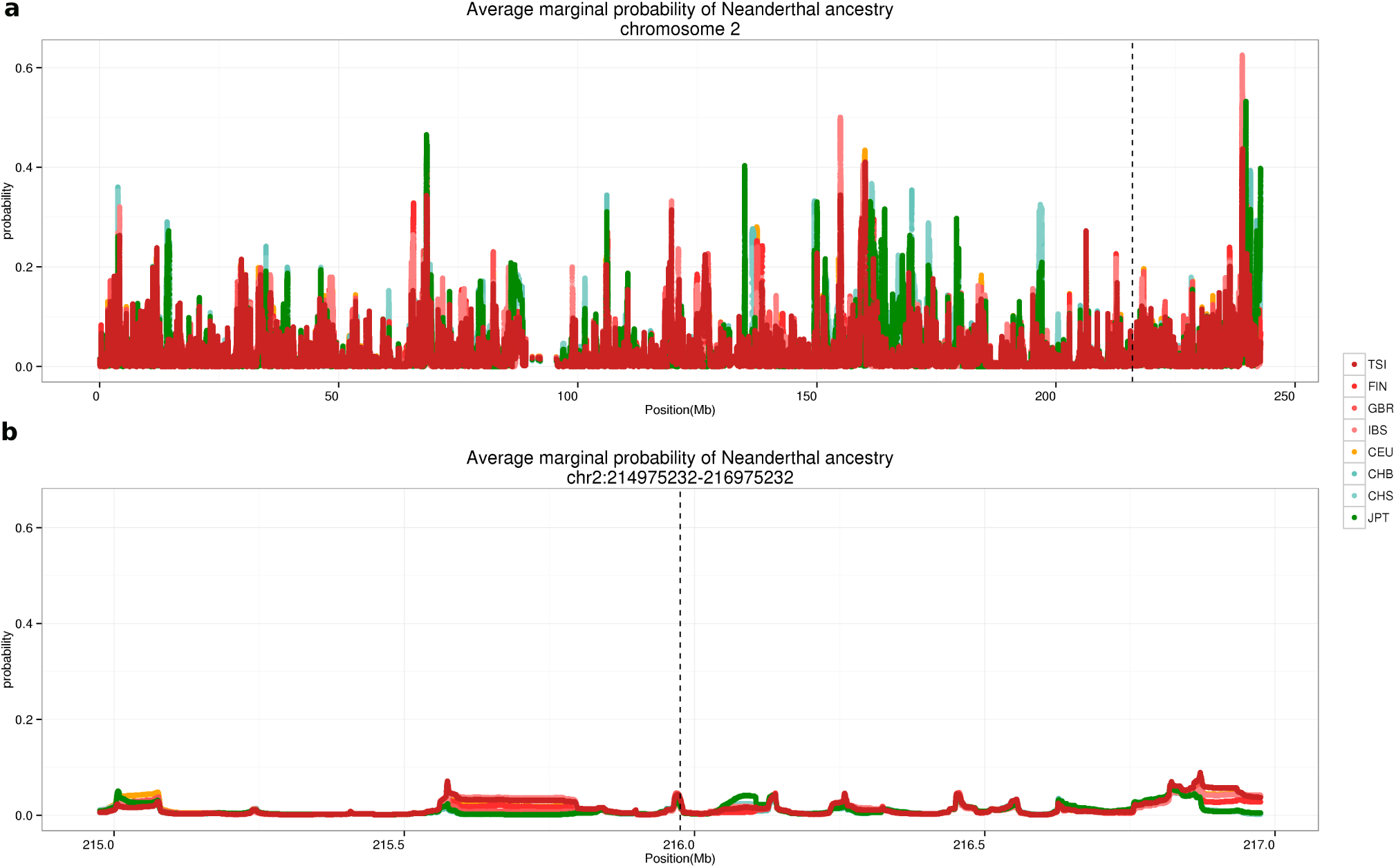
Neanderthal introgression in Sapiens at chromosome 2. Average marginal probability of introgression in Eurasian populations according to^34^. Dots (very dense) represent probability in a 100 kb region in a group of individuals from the same population for all chromosome 2 (**a**) and a 2kb region surrounding rs10180970 (**b**). The gray dashed line represent the genomic position of rs10180970.

**Figure S3.**
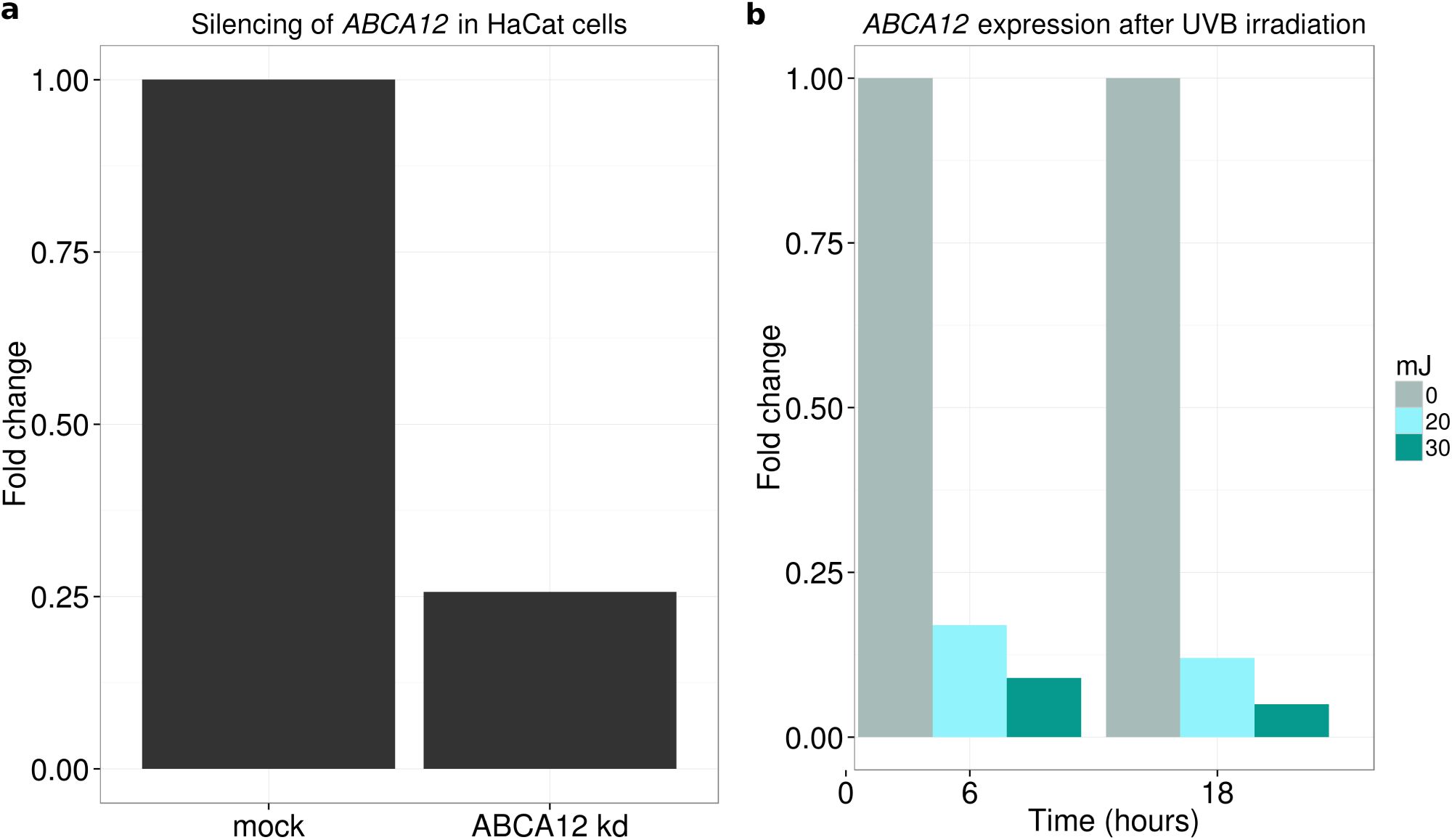
*ABCA12* expression in HaCaT cells and UVB irradiation. (**a**) *ABCA12* is expressed in HaCaT cells and can be silenced using small interfering RNA. (**b**) *ABCA12* expression is reduced after Ultraviolet B (UVB) radiation and cells do not recover even after 18 hours. T6 and T18 indicate 6 and 18 hours after UVB radiation; mJ = milliJoule.

